# Miniaturised laboratorial equipment as a solution to implement conservation genetics tools and education in West African countries with limited infrastructures: an application to the study of illegal wildlife trade in Guinea-Bissau

**DOI:** 10.1101/2024.09.30.615830

**Authors:** Maria Joana Ferreira da Silva, Ivo Colmonero-Costeira, Mohamed Djaló, Nelson Fernandes, Tomás Camará, Rui M. Sá, Tania Minhós, Angelika Kiebler, Martin Grethlein, Netta Pikkarainen, Stefan Prost

**Affiliations:** CIBIO – Centro de Investigação em Biodiversidade e Recursos Genéticos, Universidade do Porto, Campus de Vairão, Rua Padre Armando Quintas 7, 4485-661 Vairão, Portugal; BIOPOLIS Program in Genomics, Biodiversity and Land Planning, CIBIO, Campus de Vairão, 4485-661 Vairão, Portugal; ONE Organisms and Environment Division, School of Biosciences, Cardiff University, Sir Martin Evans Building, The Museum Ave, Cardiff CF10 3AX, United Kingdom; CIAS, Department of Life Sciences, Universidade de Coimbra, Coimbra, Portugal; Abu village, Ganogo island, Guinea-Bissau; Anghôr village, Orango island, Guinea-Bissau; Quitafine village, Cacine, Guinea-Bissau; Centre for Public Administration and Public Policies, Institute of Social and Political Sciences, Universidade de Lisboa, Rua Almerindo Lessa, 1300-663, Lisbon, Portugal; Centre for Research in Anthropology (CRIA-NOVA FCSH/IN2PAST), Edifício 4 - Iscte_Conhecimento e Inovação, Sala B1.130, Av Forças Armadas 40, 1649-026 Lisboa, Portugal; Anthropology Department, School of Social Sciences and Humanities, Universidade Nova de Lisboa (NOVA FCSH), Avenida de Berna, 26-C, 1069-061 Lisboa, Portugal; Ecology and Genetics Research Unit, University of Oulu, Oulu, Finland; South African National Biodiversity Institute, National Zoological Garden, Pretoria, South Africa; Central Research Laboratories, Natural History Museum Vienna, Vienna, Austria

**Keywords:** Primates, wild meat, bushmeat, mitochondrial DNA, meta-barcoding, CytB gene, environmental crime

## Abstract

Illegal wildlife trade (IWT) is considered one of the largest global illegal industries that negatively impacts biodiversity and sustainable development worldwide. DNA barcoding coupled with high-throughput sequencing has been shown to be useful in identifying taxa affected by IWT and has been routinely used during the last decades. However, for countries lacking laboratory infrastructures and sequencing units or trained staff, the application of DNA barcoding tools in conservation actions and policies is limited and dependent on slow sample export processes and molecular analyses carried out abroad. Guinea-Bissau (GB) is located on the West-African coast and has one of the lowest human development indices worldwide, while being a biodiversity hotspot facing many conservation challenges due to illegal commercial hunting, and trade in bushmeat and live individuals. Here, we explore the potential of using inexpensive and portable miniaturised laboratory equipment (MLE) to i) identify species illegally traded in GB using DNA barcoding methods and ii) to improve molecular biology and conservation genetic education and training in GB. Our overarching aim is to raise awareness of the current gap between the need to apply conservation genetic technologies in GB and the inability to do so due to a lack of laboratory infrastructures, sequencing units and opportunities for molecular biology training. We show that MLE can be a solution to accelerate the use of DNA barcoding methods to understand IWT and to train students, technicians and staff from governmental agencies dedicated to investigating environmental crimes, ultimately advancing the discipline of conservation genetics in the country.

## Introduction

The investigation of wildlife crimes, including illegal wildlife trade (IWT), is one of four main topics addressed by conservation genetics (Frankham 2019), a subdiscipline of conservation biology. IWT can be described as “an environment-related crime that involves the illegal trade, smuggling, poaching, capture or collection of endangered species, protected wildlife, such as animals and plants that are subject to harvest quotas and regulated by permits, its derivatives or sub-products” (South & Wyatt 2011).

Overexploitation of natural resources, which includes IWT, is suspected to be the main driver of biodiversity loss today (Maxwell et al. 2016). Furthermore, IWT itself has been directly linked to biodiversity loss (PBES 2019) and affects species throughout the tree of life (Fukushima et al. 2019; Scheffers et al. 2019). Taxa are sold live or dead, whole or processed, and the resulting products or specimens may be used for pharmaceutical, food, pets, ornamental or traditional medicinal purposes (Nellemann et al. 2014; Stringham et al. 2021). Moreover, IWT is negatively impacting biodiversity and society in direct and indirect ways (reviewed in Mozer & Prost 2023). More than 10 years ago, IWT was considered the fourth largest global illegal trade after narcotics, humans and forged products (World Wide Fund for Nature 2012). However, its annual value is increasing steadily and has likely increased substantially since then (Nellemann et al. 2016). Actions identified as important in halting the rise in IWT (World Wide Fund for Nature 2012; Nellemann et al. 2014) are based on a good understanding of the phenomenon at the national and international scale, which requires accurate identification of the species and populations exploited. Importantly, IWT often affects low-income countries that harbour high amounts of biodiversity, so called biodiversity hotspots (UNODC 2016).

Guinea-Bissau (GB) is a small country located on the West African coast that is formed by a mainland and the Bijagós archipelago (Fig. 1, 36,125 km2, population: 2,08 million in 2022, CIA 2024; https://stat-guinebissau.com/). GB is considered an important biodiversity hotspot and a regional stronghold for iconic and threatened species, such as leopards (*Panthera pardus*), lions (*Panthera leo*), elephants (*Loxodonta cyclopsis),* and the western chimpanzee (*Pan troglodytes verus*) (Brugiére et al. 2005; Brugiére et al. 2006; Bersacola et al. 2018; Palma et al. 2023). Six protected areas and three ecological corridors were formally delimited to manage the conservation of biodiversity (Fig. 1) covering almost 26.3% of the country’s area^1^. At the same time, GB is a low-income country, where more than 60% of the population lives below the poverty line and there is a high level of inequality in income distribution (CIA 2024, ^2^,^3^). Furthermore, it has one of the lowest human development indices worldwide^4^. The structure of the economy remained unchanged over the last two decades. The primary sector (agriculture, livestock, and fishing) has been the greatest contributor to the economy (CIA 2024). Reasons explaining a stagnation of the national economy are related to the poor diversification of products, an acute shortage of a skilled workforce due to low efficiency of the education system, and very low private investment (which is the second lowest in Africa) (World Bank 2020). Moreover, GB has been politically and militarily unstable for the last decades (*i.e*., five coup d’etat and three attempts, the latest in November 2023) (Sangreman 2019). The events of military revolt and the regular disruption of the work by public institutions and a continuous replacement of governmental agencies personnel have contributed to the degradation of the country’s infrastructures. These include the distribution system of public electricity and water and maintenance of roads, even at the capital city - Bissau, and an irregular functioning of education institutions and an insufficient health care system (Ferreira 2022; Gomes 2020). The small number of year-around transitable roads to rural areas creates gaps in the distribution of goods and contributes to a general lack of awareness of the environmental and conservation issues in the country’s capital city (Ferreira da Silva et al. 2024). Economic activities based on or able to be reconciled with the conservation of biodiversity (such are ecotourism) are considered drivers for human and economic development (https://ibapgbissau.org/areas-protegidas/).

**Figure 1.**
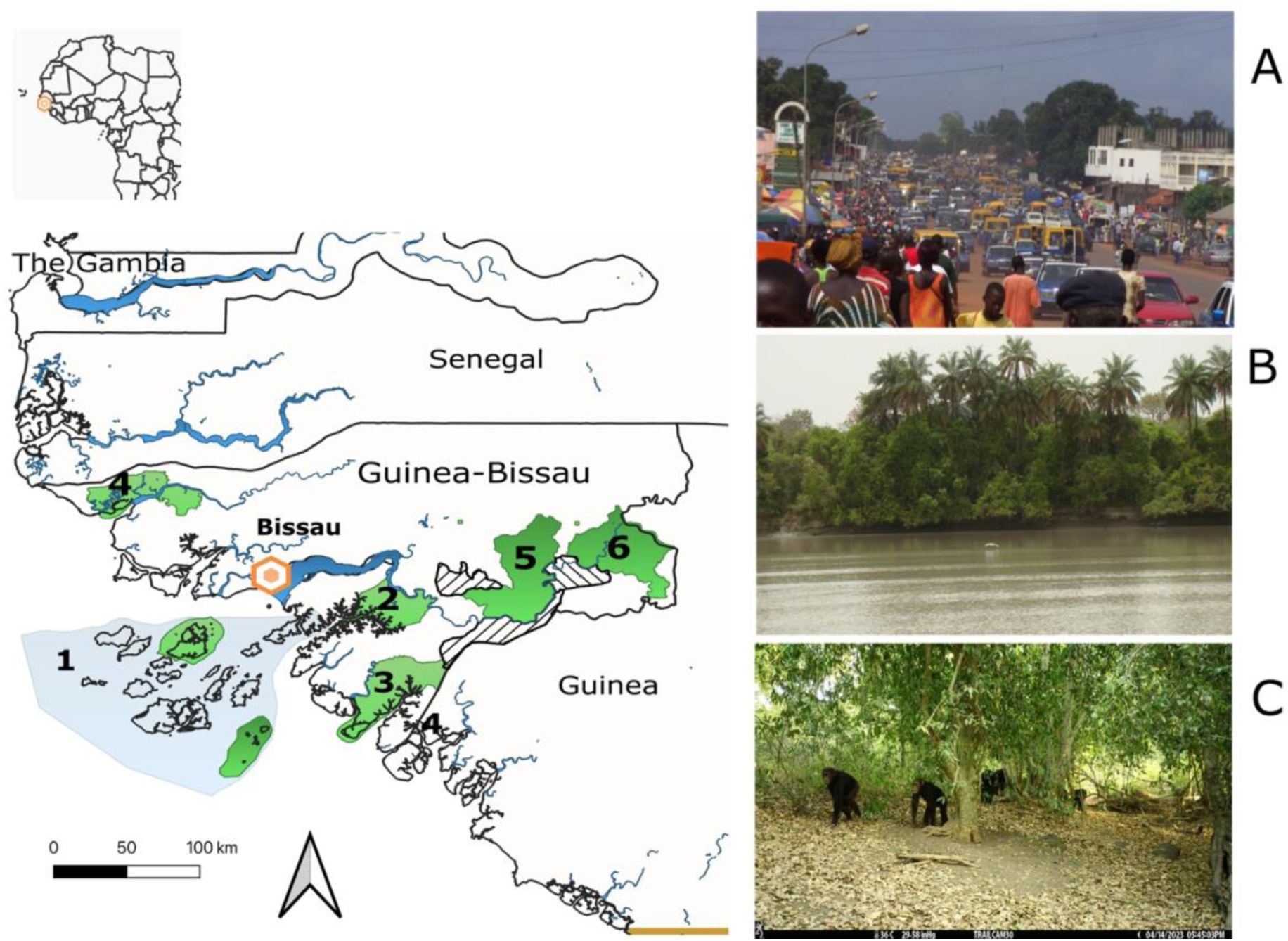
Location of Guinea-Bissau in the West African coast. The capital city – Bissau, is indicated by a hexagon. Six parks were delimited: 1 - Bolama Bijagós archipelago Biosfere Reserve, 2 – Cacheu River Mangroves Natural Park, 3 - Cufada Lagoons Natural Park, 4 – Cantanhez National Park, 5 – Dulombi National Park and 6 – Boé National Park. Three ecological corridors are indicated in black and white striped areas (information retrieved from https://ibapgbissau.org/areas-protegidas/). Legends of photos A. Bissau, the capital city (photo credits by RS); B. Corubal River, the main waterbody of the country (credits by MJFS); C. GB is a regional stronghold for the western chimpanzee (*Pan troglodytes verus*); the photo of the wild group was captured by camera trap (credits by L. Palma, TROPIBIO).

National wildlife is threatened by hunting for commercial purposes. Medium to large mammals, and non-human primates (primates hereafter) in particular, are hunted for the trade of meat, of skins and body-parts and the national and international trade of live exotic pets (Ferreira da Silva et al. 2021b; Minhós et al. 2013; Sá et al. 2012, Fig. 1). There is little information on targeted species and the profile of consumers (but see Ferreira da Silva et al. 2021b). Hunting protected wild animals to sell its meat is illegal (Decree n°21/1980, Decree n°2/2004, Decree n°3/2017) and people involved in the trade of bushmeat are reluctant to provide detailed information on activities (Costa et al. 2013; Minhós et al. 2013). Existing data are based on hunter’s qualitative descriptions of activities (e.g., Amador et al. 2014; Ferreira da Silva et al. 2021a) and on rapid surveys in trading sites and the DNA barcoding of samples collected from carcasses (Minhós et al. 2013; Ferreira da Silva et al. 2021b).

Past studies unravelled that meat from wild animals is mainly sold to private customers at meat markets and in specialised bars and restaurants, which are thought to exist across the country but that are generally hidden from outsiders (Starin 2010; Minhós et al. 2013b; Colmonero-Costeira et al. 2023). DNA barcoding confirmed the identity of species traded (Minhós et al. 2013b; Ferreira da Silva et al. 2021a). Results suggested that primate species were the most commercialised taxa, in particular the green monkey (*Chlorocebus sabeus*), the Campbell’s mona monkey (*Cercophitecus campbelli*) and the Guinea baboon (*Papio papio*) (Minhós et al. 2013b; Ferreira da Silva et al. 2021a). The trade of primates classified as threatened of extinction by the International Union for Conservation of Nature (IUCN), such the red colobus (*Piliocolobus badius temminckii*) and the black and white colobus (*Colobus polykomos*) was also detected (Minhós et al. 2013b). The study of the temporal dynamics of species consumed at commercial establishments (e.g., bars) was possible by applying meta-barcoding tools (e.g., Ferreira da Silva et al., 2021a). Results suggested that the Campbell’s mona monkey is the most consumed taxon in these contexts (Ferreira da Silva et al., 2021a).

However, the molecular identification of tissue samples collected in IWT contexts in GB were done abroad because the country lacks molecular laboratory infrastructures where DNA barcoding protocols could be carried out (Ferreira da Silva, MJ personal observation^5^). This is a common challenge faced in West Africa (Kanteh et al. 2022 REF Helmy et al. 2016; (REF). In GB specifically, the single local university teaching molecular biology and related disciplines (as conservation genetics) within the context of a degree in a biology thematic area^6^ is not equipped with molecular laboratory facilities (Sá 2019). This is because electricity is very unstable at the capital city and cuts in the supply are frequent, which would damage the laboratory equipment (R. Sá, personal communication^7^). Outside the university, at least two private labs hold chambers for DNA extractions and a PCR machine, but these have specific purposes, namely, to detect the presence of the Ebola virus^8^, or COVID-19 virus from human samples (Kanteh et al. 2022), which do not include conservation genetic applications. Moving biological samples obtained in IWT contexts from GB to a European country for molecular identification has many disadvantages. The two most important drawbacks are i) the issue of CITES exportation permits by national focal authorities is extremely slow^9^ and has delayed the production of data, which prevented the understanding of the IWT phenomena in a timely manner to act upon; ii) the staff of agencies responsible for law enforcement and conservation management (*i.e.,* Direcção Geral da Fauna e Flora, DGFF, and Instituto para a Biodiversidade e Áreas Protegidas, IBAP) and other conservation practitioners such as non-governmental agencies (e.g., CHIMBO) are not trained in genetic or genomic technologies within the country and miss important learning opportunities offered by these studies.

DNA barcoding tools have been widely and effectively used during the last decade to identify taxa affected by IWT (Staats et al. 2016). These technologies are particularly important in cases of impaired visual identification such of processed specimens and products, and for monitoring poaching and commercial trade (e.g., Eaton et al. 2010; Olayemi et al. 2011; Minhós et al. 2013b; Gaubert et al. 2015; Ferreira da Silva et al. 2021a). Another important application of these techniques is the determination of the geographic origin of confiscated specimens and hunting hotspots (e.g., chimpanzees *Pan troglodytes* spp., Fontsere et al. 2022 and elephants, *Loxodonta* spp., Wasser et al. 2015). DNA barcoding tools require the amplification by Polymerase Chain Reaction (PCR) of short (*e.g.,* less than 200-600 base pairs (bp)) and taxonomically informative DNA sequences. The most frequently targeted mitochondrial (mtDNA) region is the cytochrome c oxidase 1 (COI, Hebert et al. 2003), although other genes may be used (*e.g.*, *cytb*, 12S and 16S, Gaubert et al. 2015). The genetic information obtained from specimens’ samples is then compared to vouchers deposited in reference datasets, such as the Barcode of Life Data System (BOLD, Ratnasingham and Hebert 2007), the National Center for Biotechnology Information (NCBI) GenBank database (Benson et al. 2013) or DNA_BUSHMEAT_ (Gaubert et al. 2015), to obtain a molecular identification. The recent use of high-throughput sequencing (HTS) technologies as a replacement of the more traditional Sanger sequencing have been considered advantageous in some IWT contexts (Staats et al. 2016). HTS technologies allow multiple DNA barcode templates and samples to be sequenced in parallel. This also opened up the possibility to sequence mixed community samples and environmental samples containing DNA of tens to thousands of species, an approach that is referred to as meta-barcoding. The use of DNA and meta-barcoding have many advantages in the investigation of the IWT as they allow for i) a higher number of samples to be analysed simultaneously, which reduces the costs and time of laboratory procedures, ii) the success of analyses of samples containing highly degraded DNA is improved, namely samples collected from cooked meat or pre-processed products, and iii) multiple species can be identified in mixed samples, which are typical of samples collected from traditional Chinese medicines (Coghlan et al. 2015) or meals with different types of meat (Ferreira da Silva et al. 2021a; Staats et al. 2016).

Nevertheless, despite a recognized need to integrate molecular tools to inform conservation actions, there is a mismatch between availability of genetic technologies (and genetic data) and its application in conservation actions and policies (Garner et al. 2016; Klütsch & Laikre 2021; Bertola et al. 2024). Amongst other reasons, a routine use of genetic and genomic technologies to identify taxa in IWT may not be possible because of i) lack of knowledge on how to initiate, apply or use these tools or difficulty in interpreting and communicating results to funders and collaborators (Holderegger et al. 2020), ii) prohibitive costs to analyse a large amount of samples, iii) limited or no availability of laboratory facilities with high-throughput sequencing equipment and high-performance computing clusters, and lack of university-trained technicians able to do such lab protocols (Klütsch & Laikre 2021; in sub-Saharan Africa, Helmy et al. 2016), iv) delays in samples’ exportation to countries equipped with molecular laboratory facilities due to international regulations (e.g., Convention on International Trade in Endangered Species [CITES]) or difficulties in obtaining permits to carry out genetic sampling by foreign teams (e.g., associated with the Nagoya Protocol on Access and Benefit Sharing, https://www.cbd.int/abs/) (Bertola et al. 2024).

The use of miniaturised laboratory equipment (MLE) has the potential to resolve some of the challenges of applying conservation genetics technologies in countries lacking molecular laboratory facilities, trained personnel or dedicated-funding lines. MLE are portable and inexpensive laboratory instruments that can be used for DNA extraction, PCR and mtDNA sequencing (e.g. Pomerantz et al. 2022). MLE devices comprise benchtop centrifuges, thermocyclers, gel electrophoresis systems, magnetic racks to perform bead cleanups of PCR products and of use during library preparations steps before sequencing, and sequencing devices based in nanopore nucleic acid sequencing (ONT) such as the MinION (see Fig. 1 in Pomerantz et al. 2022). MLE allows for the molecular identification of taxa in different circumstances and contexts, including the ones in which electricity and Internet access are inconsistent. For instance, Pomerantz et al. (2018) obtained high-accuracy consensus sequences that allowed for the identification of species, including rare ones, 24h after collecting the specimens in the Ecuadorian Chocó rainforest. DNA and meta-barcoding protocols using MLE have been developed specifically for non-standard contexts. For instance, to analyse a large number of samples on-site relatively quickly and using a minimal number of MLE devices, and for analyses to be carried out by personnel with little experience in bioinformatic pipelines and using personal laptops (Pomerantz et al. 2022). A key equipment in the complete processing of DNA barcode data *in situ* is the MinION, an inexpensive and portable sequencing platform from Oxford Nanopore Technologies (ONT, UK). Using a nanopore-based technology allows the sequencer to be very small (10 × 3.2 × 2 cm and 90 grams) compared to other high-throughput sequencing platforms (https://nanoporetech.com/). It has been shown to reliably work for in situ molecular analyses (e.g., Blanco et al. 2020, Menegon et al. 2017, Pomerantz et al. 2019, Srivathsan et al. 2021), education (e.g. Salazar et al. 2020, Watsa et al. 2020) and for medical applications (e.g. Quick et al. 2018). Moreover, recently the use of ONT’s MinION sequencing platform coupled with a DNA barcoding analysis software (NGSpeciesID, Sahlin et al. 2021) has been validated for its use in wildlife forensic application (Vasijevic et al. 2021; reviewed in Ogden et al. 2021).

In this work, we aimed to discuss and explore the use of MLE and ONT’s MinIOn sequencer together with a published bioinformatic pipeline (Pomerantz et al. 2022) in the molecular identification of samples collected from illegally traded bushmeat carcasses in GB and its potential for local capacity sharing and training. Tissue samples were collected between 2010 and 2022 in different contexts – bushmeat markets, bushmeat-dedicated restaurants, and at a stop-point in the transportation route from the hunting area to the final customer. We selected samples to be representative of different situations faced by researchers and the sample’s preservation conditions. Our overarching aim is to raise awareness for the current gap between the need and inability to apply conservation genetics technologies in GB and possibly in other West African countries lacking large laboratory infrastructures and sequencing units and to propose MLE as a solution to accelerate the use of DNA barcoding-based technologies in efforts to understand IWT in these contexts and develop conservation genetics as a discipline.

## Methods

### Study area and samples collection

This study used 33 tissue samples collected as part of other studies that investigated wild meat hunting, trade and consumption in several locations in Guinea-Bissau (Minhós et al. 2013b; Ferreira da Silva et al. 2021a) (Fig. 2). Here, we analysed i) eleven samples collected in Bissau in 2010 in two markets (Minhós et al. 2013b), ii) 12 samples collected as part of a study carried out from 2015 to 2017 that monitored the consumption of wild meat at dedicated restaurants in the south of the country (Ferreira da Silva et al. 2021a) and, iii) ten tissue samples collected from an island of the Bijagós Archipelago between May and June 2022, which were obtained during monitoring the exportation of wild meat from the main seaport of the island to the urban wild meat markets in Bubaque island and the capital city, Bissau.

**Figure 2.**
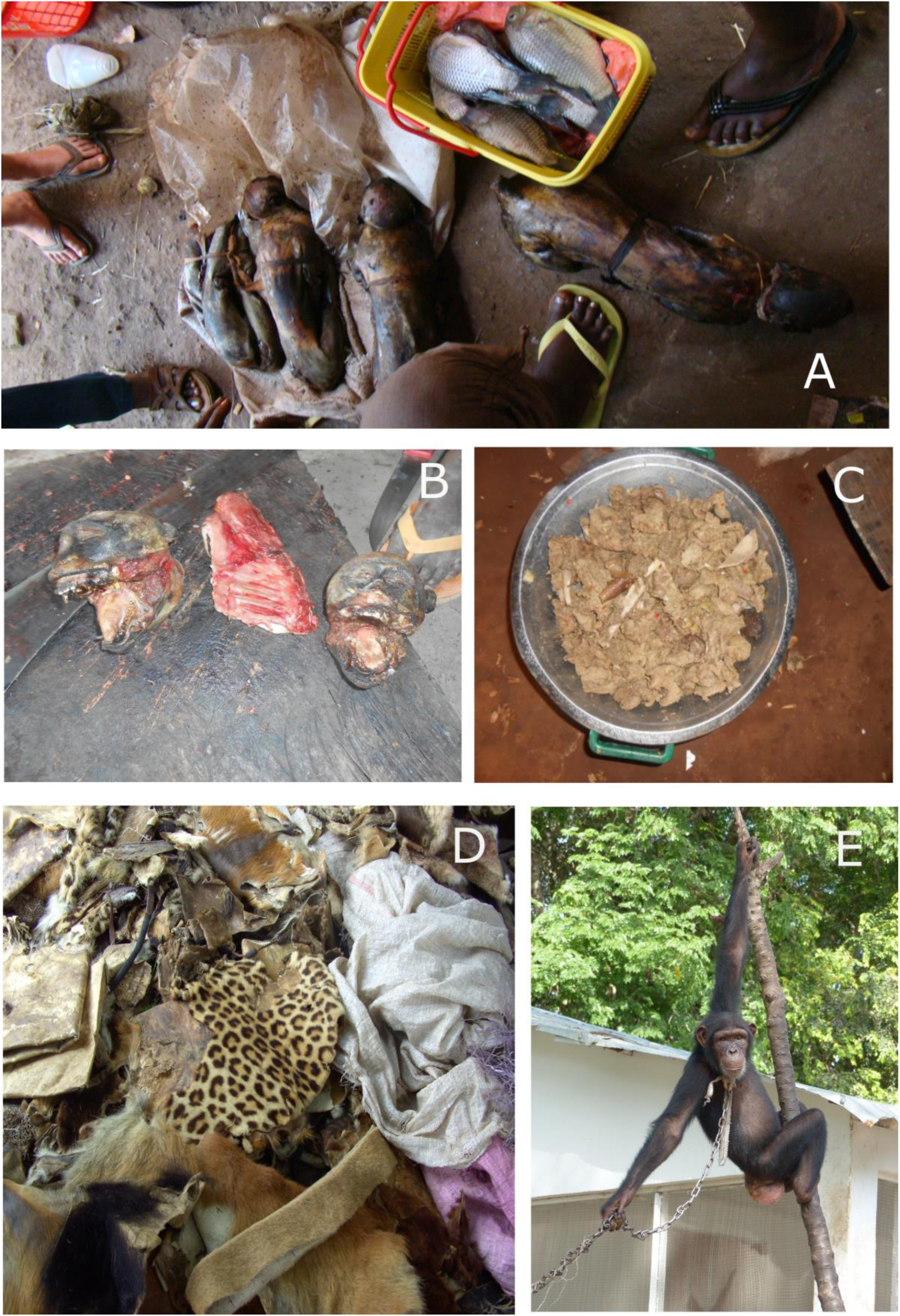
Examples of commercial hunting activities in Guinea-Bissau. Photo A. Aspect of wildmeat markets in the capital city – Bissau. Both fish and wildmeat were being traded at the same location. Carcasses of primates arrived whole, disembowelled and smoked, making the visual identification of species inaccurate (photo credits by D. Starin, 2010). B. heads of Guinea baboons (*Papio papio*) being prepared for consumption in a smaller town in the south of the country (photo credits by T. Camará). C. Aspect of a meal containing primate meat served in restaurants and bars in Guinea-Bissau (Photo by T. Camará, more information in Ferreira da Silva et al., 2021a). D. Trade of animal’s body parts in Bandim market, the largest generalist market in the capital city (credits by F. Sousa, more information in Sá et al 2012). The stall was selling skins from leopards, western chimpanzees, Guinea baboons, and possibly Red colobus monkeys. E. A photo of the chimpanzee Emily, one confiscated chimpanzee from the national trade of exotic pets.

Tissue samples were placed in provided 2 ml Eppendorf tubes containing 98% ethanol using pre-named zip-lock plastic bags and a coding scheme. We specifically collected tissue from unburnt/uncooked parts of carcasses. Samples were exported to Portugal.

### DNA extraction, amplification and sequencing

DNA was extracted using ThermoFisher’s MagMAX DNA Multi-Sample Ultra 2.0 Kit on the ThermoFisher Kingfisher Apex DNA extraction platform using default parameters. For the PCR amplification we used (1) primers designed by Gaubert et al. (2015) for the identification of bushmeat samples that amplify the first 402 bp of cytochrome b (*cyt b*) and a 390 bp of the 12S region, and (2) approximately 250 bp of the 16S rRNA gene using primers from Vences et al. (2016) that were designed to amplify vertebrate DNA from environmental DNA samples. Fragments were amplified by PCR in 30ul total volume reaction, using 15 ul of Qiagen Multiplex PCR Mix (Qiagen, GER), 0.6 uM of forward and reverse primers and 1ul of DNA. Thermal cycling was performed in a T100 Thermal Cycler (BioRad). Cycling conditions consisted of 94°C for 3 min, followed by 35 cycles at 92 °C for 30’’, the primers specific annealing temperature for 30 s (CytB – 48°C, 12S – 56°C and 16S – 59°C) and 72°C for 30 s, followed by a final extension of 72°C for 15 min. A template-free PCR was included in each amplification to control for potential contamination. Amplification success was checked on 2% agarose gels.

The sequencing was carried out on the ONT’s MinION Mk1C platform using the native barcoding kit (ONT, SQK-NBD114.96) according to the manufacturer’s manual with minimal exceptions. In brief, the ends of the DNA amplicons were first prepared for index ligation using the NEBNext Ultra II End Repair / dA-tailing module (NEB, USA), here we extended the reaction time to 30 min at 20C and 30 min at 65C, then individual indices were aligned using the NEB Blunt/TA Ligase Master Mix (NEB, USA), cleaned with AMPure XP Beads (Beckman Coulter, USA) and then pooled in equal ratios. Sequencing adapters were ligated to the indices using the NEBNext Quick Ligation Module (NEB, USA) and the reaction cleaned using AMPure XP Beads (Beckman Coulter, USA) with ONT’s Short Fragment Buffer (SFB). The final library was quantified using the Qubit 4 Fluorometer and about 200ng loaded onto an ONT Flongle sequencing flowcell.

### Bioinformatic pipelines and assignment of samples

We followed the bioinformatic pipeline described in Pomerantz et al. 2022. First the raw pod5 files were converted to the fastq format and demultiplexed using Dorado 0.5.0 (https://github.com/nanoporetech/dorado). Next, the number of reads and their quality were assessed using NanoPlot (https://github.com/wdecoster/NanoPlot) and we only retained reads with a Phred score of at least 17 with NanoFilt (https://github.com/wdecoster/nanofilt). Priming sites were removed from the reads using cutadapt (https://github.com/marcelm/cutadapt/; Martin 2011) and the consensus sequences reconstructed using NGSpeciesID (Sahlin et al. 2021). We carried out consensus polishing within NGSpeciesID using ONT’s Medaka software (https://github.com/nanoporetech/medaka). To identify the sample to the species level, we compared the sequences to available data on NCBI (http://www.ncbi.nlm.nih.gov/) using Nucleotide BLAST (Boratyn et al. 2013).

### Ethical guidelines

This research involved volunteer participants who were informed of the procedures and possible risks involved. Oral informed consent to participate was obtained before beginning research. Participants understood they could stop participating in the study at any time. The restaurant’s clients were aware that restaurants serve wild meat, and our team did not interact with them. Confidentiality of the information provided/observed, and anonymity of identity and location of establishments was given to participants. The identity of participants was not made available to law enforcers. Invasive samples of endangered species were always obtained freely. Ethical guidelines were approved by the ethics committee of the University of Porto, Portugal (REPORT N° 127 /CEUP/2022) and Cardiff University, UK (SREC 22 11-01). Guinea-Bissau authorities approved the use of tissue samples for research, the timeline and the research plan and were acknowledged in research dissemination activities. IBAP (Institute for Biodiversity and Protected Areas) and DGFF (Direção Geral de Florestas e Fauna, CITES focal point) authorized the study, the collection of samples and issued permits to transport samples to Portugal, where lab work was carried out. CITES samples importation authorization was provided by Instituto para a Conservação da Natureza e Florestas (ICNF, Portugal) (*Cercopithecus petaurista* - N.°18PTLX00592I and 23PTLX00512I, *Cercopithecus campbelli* - N.°18PTLX00590I, *Erythrocebus patas* - 8PTLX00589I, *Chlorocebus sabaeus* - N.°/No 18PTLX00586I, *Papio papio* - 18PTLX00585I). Direção Geral de Veterinária (DGV) issued health and veterinary permits to import tissue samples to Portugal. Tissue samples were transported in compliance with DOT and IATA triple packaging requirements. The research was made entirely available to national organizations. The research complied with health and safety protocols in place at the laboratory facilities to reduce harm to the environment and to research staff. The researcher did not handle or harmed animals and only used specimens’ already diseased.

## Results

Whole genome DNA was successfully extracted from the tissue samples; PCR and sequencing procedures worked equally well for all DNA extracts.

The 99 mtDNA sequences obtained (three markers per sample: CytB, 12S and 16S) showed high percentages of identity to vouchers deposited in GenBank. In all but one case the three genetic markers agreed on the species identification (Table 1). In this case (sample 21, see Table 1), the 16S and the CytB identified the sample as *Cercopithecus campbelli*, while the 12S marker identified it as *Cercopithecus mona*. However, the sample (GenBank accession: <u>JQ256980.1</u>; voucher accession: NHMUK:ZD.1956.276) originated from Ghana, where both species occur.

**Table 1.** Results from molecular identification using nanopore-based nucleic acid sequencing (Oxford Nanopore Technologies (ONT) and a published bioinformatic pipeline (Pomerantz et al. 2022). ID of tissue samples collected from illegally traded wildmeat carcasses in Guinea-Bissau, previous information on species ID and Species identification carried out in this work.

| Samples ID | Previous information on species ID | Species identification in this work |  |  |
| --- | --- | --- | --- | --- |
|  |  | *cytb* | 12S | 16S |
| 224 | *Suspected*<br>*Cercopithecus petaurista* | *C. petaurista* | *C. petaurista* | *C. petaurista* |
| 225 |  | *C. petaurista* | *C. petaurista* | *C. petaurista* |
| 276 |  | *C. petaurista* | *C. petaurista* | *C. petaurista* |
| 278 |  | *C. petaurista* | *C. petaurista* | *C. petaurista* |
| 277 |  | *C. petaurista* | *C. petaurista* | *C. petaurista* |
| 280 |  | *C. petaurista* | *C. petaurista* | *C. petaurista* |
| 258 |  | *C. petaurista* | *C. petaurista* | *C. petaurista* |
| 256 |  | *C. petaurista* | *C. petaurista* | *C. petaurista* |
| 254 |  | *C. petaurista* | *C. petaurista* | *C. petaurista* |
| 255 |  | *C. petaurista* | *C. petaurista* | *C. petaurista* |
| B24 | *Papio papio* | *P. papio* | *P. papio* | *P. papio* |
| B115 | *Papio papio* | *P. papio* | *P. papio* | *P. papio* |
| B197 | *Papio papio* | *P. papio* | *P. papio* | *P. papio* |
| B155 | *Papio papio* | *P. papio* | *P. papio* | *P. papio* |
| B20 | *Erythrocebus patas* | *E. patas* | *E. patas* | *E. patas* |
| B126 | *Papio papio* | *P. papio* | *P. papio* | *P. papio* |
| B415 | *Papio papio* | *P. papio* | *P. papio* | *P. papio* |
| B127 | *Papio papio* | *P. papio* | *P. papio* | *P. papio* |
| B120 | *Papio papio* | *P. papio* | *P. papio* | *P. papio* |
| B119 | *Papio papio* | *P. papio* | *P. papio* | *P. papio* |
| B17 | not assigned | *P. papio* | *P. papio* | *P. papio* |
| B117 |  | *P. papio* | *P. papio* | *P. papio* |
| 205 |  | *Piliocolobus badius* | *Piliocolobus badius* | *Piliocolobus badius* |
| 209 |  | *Erythrocebus patas* | *Erythrocebus patas* | *Erythrocebus patas* |
| 206 |  | *P. papio* | *P. papio* | *P. papio* |
| 208 |  | *Chlorocebus sabaeus* | *Chlorocebus sabaeus* | *Chlorocebus sabaeus* |
| 203 |  | *Cercopithecus campbelli* | *Cercopithecus campbelli* | *Cercopithecus campbelli* |
| 202 |  | *Chlorocebus sabaeus* | *Chlorocebus sabaeus* | *Chlorocebus sabaeus* |
| 20 |  | *P. papio* | *P. papio* | *P. papio* |
| 201 |  | *P. papio* | *P. papio* | *P. papio* |
| 204 |  | *Piliocolobus badius* | *Piliocolobus badius* | *Piliocolobus badius* |
| 21 |  | *Cercopithecus campbelli* | *Cercopithecus mona* | *Cercopithecus campbelli* |
| 1 |  | *Chlorocebus sabaeus* | *Chlorocebus sabaeus* | *Chlorocebus sabaeus* |

## Discussion

Here, we discuss and explore the use of MLE and ONT’s MinIOn sequencer together with a published bioinformatic pipeline (Pomerantz et al. 2022) in the molecular identification of samples collected from illegally traded wild meat carcasses in GB and demonstrate its potential for the advance of local-capacity building and education in conservation genetics.

### Wild meat trade and its ecological and societal impacts in Guinea-Bissau

Commercial hunting to supply the illegal trade of wild meat, live individuals and body-parts used in traditional medicine (Ferreira da Silva et al. 2021a, Sá et al. 2012) are probably the most significant threats affecting wildlife in Guinea-Bissau. Although hunting and trade of the carcasses of primates are illegal, meat is traded in numerous urban meat markets and presumably in specialised restaurants and bars, which are thought to be scattered in urban and semi-rural areas (Minhós et al. 2013; Ferreira da Silva et al. 2021a). At these establishments, the consumption of wild meat is associated with alcoholic drinks (Minhós et al. 2013). At restaurants in the capital city in 2010, the typical dish contains four pieces of meat cooked in a stew and cost 2.60 US$ (Minhós et al. 2013). This dish was considered a luxury considering that in the country most of the population live with less than 2 US$ a day (Starin 2010). In contrast, at a town located in the south of the country, wild meat is sold by the piece at 0.10 US$ (Ferreira da Silva et al. 2021a), which suggests wide accessibility by local communities. Although the actors involved in the trade are reluctant to share information with outsiders, these establishments announce the commercialization of wild meat outside the premises and through social media which suggests that the owners are not afraid of penalties related to their activities or are unaware of the illegality of the trade.

Hunting towards wildlife may be unsustainable in GB. A minimum of 1,500 specimens was extrapolated to be traded in two urban markets in Bissau during the dry season (Minhós et al. 2013). These high numbers of hunted animals may be leading to significant demographic changes in wild populations. Genetic evidence for recent population declines was found for populations within parks (e.g., colobus monkeys in CNP, Minhós et al., 2016, chimpanzees in CNP and CLNP, Ferreira da Silva et al. in review, and petaurista monkeys in the islands, Colmonero-Costeira et al., in review) and for recent changes in gene flow patterns (for the Guinea baboon, Ferreira da Silva et al. 2014; 2018, and colobus monkeys, Minhós et al. 2013). Modifications are attributable to the joint effects of hunting and habitat loss and fragmentation. Moreover, primates at CNP display behaviours which are different from what have been described elsewhere likely as an adaptation to recent human-induced alterations in the habitats (Minhós and Ferreira da Silva 2020). These modified behaviours include i) a preference for human-cultivated foods while wild fruits are available in the forest by the western chimpanzee (Bessa et al. 2015), which further increase the contact between chimpanzees and human communities and, ii) the use of unusual sleeping sites by the Guinea baboons, such as mangrove trees, to avoid being reached by hunters during the night and reduced long-distance vocalisations (Minhós and Ferreira da Silva 2020).

### Monkey species affected by bushmeat trade in GB

An accurate description and characterization of conservation threats is the first step to improve conservation management. The use of DNA barcoding tools was key to advance the knowledge of the species affected by IWT in GB. Before the first study using DNA barcoding tools was conducted, it was thought that the species most frequently targeted for the wild meat trade in Guinea-Bissau were the Guinea baboon followed by the green monkey (*C. sabaeus*) and Colobus monkeys (*Piliocolobus badius temminckii* and *Colobus polykomos*) (Casanova and Sousa, 2007; Cá, 2008). By investigating the wild meat trade in 2010 at two urban markets in Bissau using a COI gene fragment and sanger sequencing, molecular evidence that six out of the 10 primate species in the country were traded in these contexts was provided (Minhós et al 2013). The previous suspicion of widespread hunting practices towards primates (Gippoliti and Dell’Omo, 2003; Casanova and Sousa, 2007) was confirmed, however the species-specific contribution to the trade was unexpected. *C. sabaeus* and the Campell’s monkey (*C. campbelli*) were the most frequently traded species (32.9 % and 29.7 %, respectively), despite these species having a relatively smaller body size when compared to the Guinea baboon (Rowe 2016) and presumably representing a lower financial return in the trade (Ferreira da Silva et al. 2021b). Also, colobus monkeys were not as frequently traded as initially thought (Minhós et al 2013). We also found that the use of a DNA barcoding approach was very effective in clarifying misidentifications by vendors. The non-use of DNA barcoding would lead to a wrong estimate of species frequencies in trade and redirect conservation efforts to less affected species while neglecting the most threatened ones (Minhós et al., 2013). At the trade sites, carcasses were traded smoked and disembowelled, and were generally deprived of external features which could be used for visual identification (Fig. 2). Also, there were no scales to quantify the weight of carcasses, and the price of specimens was established based on the body size, assessed visually by vendors (Minhós et al., 2013). The use of molecular tools showed that vendors misidentified *C. campbelli* and *C. sabaeus* frequently because the two species have similar body sizes (Rowe 2016) and the common names in Guinea-Bissau creole can be used interchangeably (Minhós et al. 2013). Without the use of DNA barcoding tools, *C. sabaeus* would be considered the single most traded species and most importantly, the trade of *C. campbelli* would be ignored. Finally, we were able to extrapolate for the first time a minimum amount of primate carcasses traded in Bissau and found that the estimated figures are similar to the ones described for other larger West African capitals (Minhós et al 2013).

Nevertheless, other than the studies conducted at two urban markets in Bissau (focussed on primates only, Minhós et al 2013) and in dedicated restaurants located in the south of the country (Ferreira da Silva et al 2021a), there is very limited knowledge on which and how many wild species are targeted for consumption. The number of carcasses, their prices, temporal changes in consumption and the geographic origin of specimens is unknown. Nevertheless, this information is key for effective law enforcement and to design conservation actions for changing behaviour. Carrying out molecular identification within the country would allow for faster results. However, the application of molecular biology techniques is limited given the limited financial and laboratory resources in Guinea-Bissau.

### Molecular biology and conservation genetics in Guinea-Bissau

With the civil war that began on June 7, 1998, most of the already limited infrastructure that had existed until then was destroyed or severely affected, including hospitals (e.g. “3 de Agosto” Hospital), educational and research establishments (e.g. National Institute of Studies and Research-INEP) and even the National Archives. In the aftermath of the war, and in the context of bilateral and multilateral cooperation agreements, education and health in GB became dependent on the generosity of its international partners. The teaching of biology gained new emphasis due to cooperation programs with mainly Portuguese NGOs, such as the Faith and Cooperation Foundation (FEC), which implemented a Program to Support the Reform of the Education System in Guinea-Bissau (2016-2020) based on programmatic initiatives that preceded it, which included the teaching of biology, although only at a theoretical level. One of the main challenges that persists in Guinea-Bissau is not only the lack of a network of libraries and adequate textbooks, but also the necessary teacher-training in the biology domain. In 2014 the country’s first university department dedicated to Environmental Sciences was established, as well as the first-degree course in Marine and Environmental Sciences promoted by a private university and thus not dependent on funds from the state budget, with autonomous and independent management that has able to maintain stability in the face of a troubled social and political context. With the establishment of this department and this course also came the need to set up two laboratories dedicated to experimental teaching: one for chemistry and the other for biology. Both were equipped with basic material, but they did not include equipment dedicated to molecular biology due to constraints in accessing materials as well as the high prices practiced in the sub-region. There is a great demand from Guinean university students for a more empirical and laboratory-based teaching pedagogical model. Two science fairs open to the general public have been held at the national level, with a widespread consensus that if proper investment is made in the training of young people and if they are provided with access to materials and equipment, the transformative power of education will surely yield its fruits not only in terms of social elevation but also in changing attitudes towards environment and conservation.

### Contribution to reducing barriers to equitable access and application

Here we tested the usability of miniaturised laboratory equipment for the successful identification of wild meat samples and explored its use for promoting molecular biology and training of conservation genetics methods in Guinea-Bissau. The country is considered low-income, where a large number of people live in poverty. Relatively inexpensive equipment such as MLE could help achieve several United Nations Sustainable Development Goals (https://sdgs.un.org/, SDGs). For example, it can greatly support investigations into the illegal bushmeat trade and thus support SDG15 (to aid in conservation of biodiversity) and SDG16 (by supporting the development of efficient legal prosecution of wildlife crimes) or be used in classroom environments for education and thereby supporting SDG4 (by providing quality tertiary education). The reliable use of MLE for DNA barcoding has been shown in many different contexts, including *in situ* molecular analyses (e.g., Blanco et al. 2020, Pomerantz et al. 2019; Srivathsan et al. 2021), education (e.g., Salazar et al. 2020, Watsa et al. 2020) and for medical applications (e.g., Quick et al. 2018). Furthermore, ONT’s inexpensive and portable MinION sequencer has previously been validated for its use in DNA barcoding for wildlife forensic casework (Vasiljevic, et al. 2021). This makes MLE a promising alternative to standard and often expensive molecular biology equipment.

## Acknowledgments

We acknowledge the Guinea-Bissau governmental agency Instituto de Biodiversidade e Áreas Protegidas (IBAP), in particular to the director Dr. Aissa Regalla and former director Dr. Justino Biai, and to the directors of protected areas and staff members Dr. Abilio Said, Dr. Augusto Cá, Dr. Joãozinho Mané and Dr. Sadjo Danfa for fieldwork and sampling permits and to Abel Vieira, Iaia, Benjamim, Bemba for the support in fieldwork logistics. We would like to acknowledge the role of Direcção Geral de Florestas e Fauna (DGFF) and CITES focal person in acquiring the sample export permit. We are grateful to the work of the research assistants and guides Sadjo Camará, Mamadu Soares, Mamadu Turé, Idrissa Camará, to the NGO CHIMBO for logistical support to carry out fieldwork in the Boé region and to I. Espinosa and H. Foito for logistical support in Bissau. This research was carried out as part of the PRIMACTION project and was partly funded by the Born Free Foundation, Chester Zoo Conservation Fund, Primate Conservation Incorporated, Mohamed Bin Zayed (Project 232533027) and by the Portuguese private companies - CAROSI, Cápsulas do Norte, Camarc, JA-Rolhas e Cápsulas. Fieldwork in 2022 and 2023 was partly funded by the project NORTE-01-0145-FEDER-000046, supported by Norte Portugal Regional Operational Programme (NORTE2020), under the PORTUGAL 2020 Partnership Agreement, through the European Regional Development Fund (ERDF) and by the Genetic Society (UK) and Primate Conservation Incorporated (PCI#1694). This work is part of the Biodiverse Anthropocenes research project, supported by the University of Oulu and the Academy of Finland PROFI6 funding (2021-2026), project number 336449. MJFS worked under a research contract funded by the Portuguese Foundation for Science and Technology (Fundação para a Ciência e Tecnologia, FCT) (https://doi.org/10.54499/CEECIND/01937/2017/CP1423/CT0010). ICC was supported by a FCT PhD scholarship (https://doi.org/10.54499/SFRH/BD/146509/2019). RMS was supported by Portuguese national funds through FCT under project UID/00713/2020 and a contract also funded by FCT (https://doi.org/10.54499/DL57/2016/CP1456/CT0002).

## Statements & Declarations

### Competing Interests

The authors have no relevant financial or non-financial interests to disclose.

## Author Contributions

### CRediT Author Statement

**Maria Joana Ferreira da Silva** Conceptualization, Methodology, Validation, Formal analysis, Investigation, Supervision, Project administration, Funding acquisition, Writing - Original Draft, Writing - Review & Editing. **Ivo Colmonero-Costeira** - Conceptualization, Methodology, Investigation, Visualization, Funding acquisition, Writing - Original Draft, Writing - Review & Editing. **Mamadu Djaló** Investigation, Writing - Review & Editing. **Nelson Fernandes** Investigation, Writing - Review & Editing. **Tomás Camará** Investigation, Writing - Review & Editing. **Tania Minhós** Investigation, Writing - Review & Editing. **Angelika Kiebler** Methodology, Writing - Original Draft, Writing - Review & Editing**. Martin Grethlei** Methodology, Writing - Original Draft, Writing - Review & Editing. **Netta Pikkarainen** Methodology, Writing - Original Draft, Writing - Review & Editing. **Rui Sá** Writing - Original Draft, Writing - Review & Editing. **Stefan Prost** - Conceptualization, Methodology, Resources, Data Curation, Funding acquisition, Writing - Original Draft, Writing - Review & Editing

## Data Availability

Datasets generated during and/or analysed during the current study are available in the GenBank repository *[PERSISTENT LINK TO DATASETS* and are available from the corresponding author on reasonable request.

## Footnotes

1 https://ibapgbissau.org/areas-protegidas/

2 https://web.archive.org/web/20121111054424/

3 http://data.worldbank.org/country/guinea-bissau#

4 https://hdr.undp.org/system/files/documents/global-report-document/hdr2023-24reporten.pdf

5 MJFS has two-decades experience in carrying out conservation genetic studies in GB

6 “Sea and environmental sciences licentiate degree*”* (at Lusófona University-Bissau, 8 semesters)

7 R. Sá was the founder director of the degree in a biology thematic area

8 https://www.insa.min-saude.pt/visita-guiada-ao-laboratorio-portugues-para-o-ebola-na-guine-bissau/

9 In the studies carried-out by MJFS and ICC, the time from requesting the export CITES permits and obtaining CITES import permits from Portuguese authorities took on average two years and caused a significant delay in attaining the results.

## Notes

### Competing Interest Statement

The authors have declared no competing interest.

